# Brain state-dependent cortico-hippocampal network dynamics are modulated by postnatal stimuli

**DOI:** 10.1101/2020.12.25.424297

**Authors:** Yoshiaki Shinohara, Shinnosuke Koketsu, Hajime Hirase, Takatoshi Ueki

## Abstract

Neurons in the cerebral cortex and hippocampus discharge synchronously in a brain state-dependent manner to transfer information. Published studies have highlighted the temporal coordination of neuronal activities between the hippocampus and a cortical area, however, how the spatial extent of cortex activity relates to hippocampal activity remains largely unknown. We imaged macroscopic cortical activity while recording hippocampal local field potentials in unanesthetized GCaMP-expressing transgenic mice. We found that cortical activity elevates before and after hippocampal sharp wave ripples (SWR). SWR-associated cortical activities occurred predominantly in vision-related regions including visual, retrosplenial and prefrontal cortex. While pre-SWR cortical activities were frequently observed in awake and sleep states, post-SWR cortical activity decreased significantly in sleep. During hippocampal theta oscillation states, phase-locked oscillations of calcium activity was observed throughout the entire cortex state. Environmental effects on cortico-hippocampal dynamics were also assessed by comparing mice reared in an enriched environment (ENR) or under isolated conditions (ISO). In both SWR and theta oscillations, mice reared in an isolated condition exhibited clearer brain state-dependent dynamics than those reared in an enriched environment. Our data demonstrate that the cortex and hippocampus exhibit heterogeneous activity patterns that characterize brain states, and postnatal experience plays a significant role in modulating these patterns.

**Significant Statement:** The hippocampus is a center for memory formation. However, the memory formed in the hippocampus is not stored forever, but gradually transferred into the cerebral cortex. As an underlying mechanism, phase-locked synchronized activities between the cortex and hippocampus has been hypothesized. However, spatio-temporal dynamics between hippocampus and whole cortical areas remained mostly unknown. We measured cortical calcium activities with hippocampal electroencephalogram (EEG) simultaneously, and found that the activities of widespread cortical areas are temporally associated with hippocampal EEG. The cortico-hippocampal dynamics is primarily regulated by animal awake/sleep state. Even if similar EEG patters were observed, temporal dynamics between the cortex and hippocampus exhibit distinct patterns between awake and sleep period. In addition, animals’ postnatal experience modulates the dynamics.

## Introduction

The sleep-wake cycle is a fundamental rhythm of the brain. Ample evidence from psychological and behavioral studies shows that experience acquired during wakefulness is consolidated to long-term memory during sleep (Squire and Alvarez, 1995; Born and Wilhelm, 2012). According to the “two-stage model” of memory, experience is first stored in a labile representation by the cortico-hippocampal interface and later is reactivated in the hippocampus to be consolidated in to the higher-order cortex (Buzsaki, 1989; Frankland and Bontempi, 2005; Mitra et al., 2016). Two distinct hippocampal EEG patterns, theta oscillations (8-15 Hz) and large irregular activity (LIA) (Buzsaki, 1989; Pavlides and Winson, 1989; Wilson and McNaughton, 1994; Mizuseki and Miyawaki, 2017), have been linked to the encoding (online) and consolidating (offline) stages of memory formation, respectively. In particular, synchronized rhythmic cortical activity with hippocampal theta oscillations has been implicated in working memory and spatial navigation (Siapas and Wilson, 1998; Jones and Wilson, 2005; Siapas et al., 2005). Moreover, sharp wave ripples (SWRs, ~200 Hz) during LIA have been proposed to integrate hippocampal output with activities of various cortical areas including the prefrontal (Wierzynski et al., 2009) and retrosplenial (RS) (Battaglia et al., 2004), and visual cortex (Norman et al., 2019).

Theta and LIA states occur both in sleep and wakefulness. Theta states dominate during attentive behavior and REM sleep while LIA represents consummatory behavior and slow-wave sleep (Buzsaki, 1989; Foster and Wilson, 2006; Carr et al., 2011; Mizuseki and Miyawaki, 2017). A number of electrophysiology studies have been conducted to find the temporal relationship between the hippocampus and particular cortical regions (Mizuseki et al., 2011; Tang et al., 2017). While these studied have described distinct brain state-dependent temporal coordination of cortico-hippocampal activity, the spatial and temporal extent to which hippocampal activity affects the entire cerebral cortex remains elusive.

The postnatal environment strongly influences the animal’s brain function and behavior. For instance, animals reared in an enriched environment (ENR) develops higher cognitive abilities while isolated and impoverished (ISO) rearing induces anxiety-like behavior (van Praag et al., 2000; Medendorp et al., 2018). In the cortex, ENR rearing strengthens the synchrony of local field potential (LFP) patterns across multiple areas (Mainardi et al., 2014). Both cortical and hippocampal principal neurons increase their morphological complexity by ENR (Volkmar and Greenough, 1972; Moser et al., 1994). We previously reported that theta-associated gamma oscillations are facilitated in ENR animals compared to ISO animals (Shinohara et al., 2013; Hirase and Shinohara, 2014). In spite of detailed documentation of neural circuit alterations after ENR or ISO in individual brain regions, little is known how experience sculpts cortico-hippocampal network dynamics.

Here we investigated how spatio-temporal activity between cortex and hippocampus is orchestrated in distinct behavioral states by combined hippocampal LFP recording and cortex-wide Ca^2+^ imaging. We used G-CaMP7 expressing transgenic mice (Ohkura et al., 2012) that allowed monitoring of neural activity through the skull (Monai et al., 2016). This configuration enabled the assessment of cortical activity before and after SWRs as well as the determination of hippocampal theta-modulated cortical activity. We also found a distinct cortical activity pattern that are peculiar to SWRs that occur during sleep. Intriguingly, such cortico-hippocampal activity coupling is enhanced in ISO reared mice.

## Materials and Methods

### Animals

G7NG817 mice (Monai et al., 2016) were weaned at P19 and reared for 4-4.5 weeks prior to imaging (Figure 1A). Mice were reared under either ISO or ENR conditions. For ISO, mice were caged individually after weaning and raised in standard cages (length 32 cm, width 22 cm, height 13.5 cm). For ENR, 5–8 male littermates were housed together in a larger cage (length 44 cm, width 27 cm, height 18.7 cm) with a ladder, running-wheels, tunnels and toys, the location of which were changed every 5 days. Both rearing environments had a 12h/12h light/dark cycle, and water and food were given ad libitum. Prior to an imaging session, mice were anesthetized with ketamine-xylazine, an aluminum head plate with an imaging window was fixed to the mouse skull with dental cement, and a 16-channel silicon probe was implanted in the left CA1 region of the hippocampus. The cranial window and implanted silicon probe were protected from scratching or other physical damage by a plastic cover with aluminum taping. After surgery, mice were returned to their designated cages (i.e., ISO or ENR), and a habituation procedure for the head-restrained imaging/recording apparatus was performed every other day for the 7-to 10-day recovery period. All procedures involving animal care, surgery and sample preparation were approved by the Animal Experimental Committee of Nagoya City University, and performed in accordance with the guidelines of the Animal Experimental Committee of Nagoya City University.

**Figure 1.**
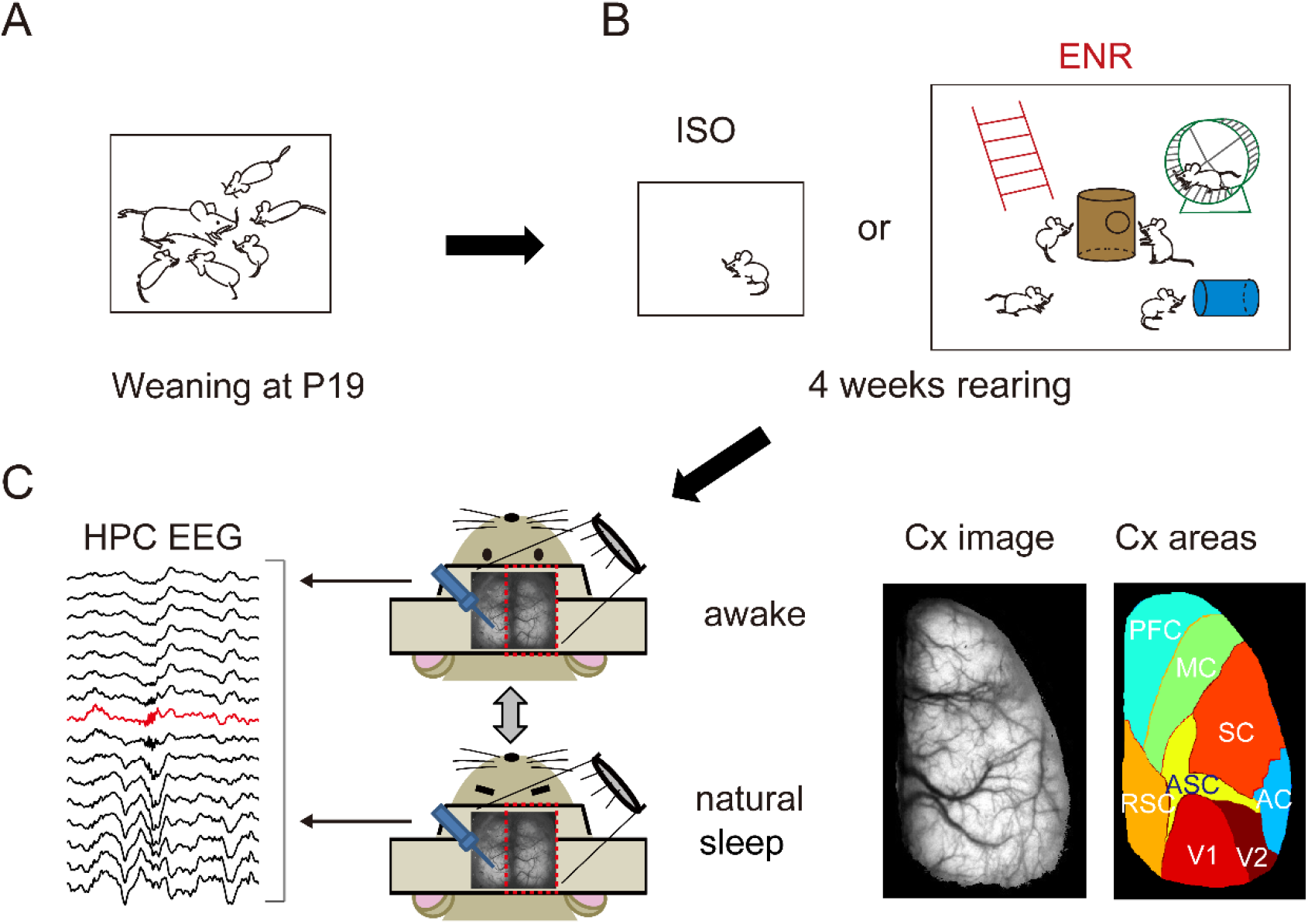
Experimental design. (A) G7NG817 mice were weaned at P19. (B) Thereafter, one male mouse is reared alone (ISO condition; left column) or 5-8 male mice were reared in a large cage with toys and wheels (ENR condition; right column) for 4 weeks. (C) Mice were restrained by a headstage (middle panel), and EEG (left panel) and imaging data (right panel) were obtained. Left panel: Representative EEG data from CA1 area in the hippocampus (HPC EEG) obtained from the left hemisphere (indicated by arrows). The red trace indicates the EEG recorded by the electrode in center of the *stratum pyramidale*. Middle panel: Animal states were monitored during the experiments and data from awake and natural sleep were analyzed separately. Right panel: Representative image data obtained from the cortex contralateral to electrode insertion (Cx image; areas surrounded by red botted rectangles in the middle panel). The cortical areas were subdivided into 8 areas (Cx areas) and analyzed further; prefrontal cortex (PFC), motor cortex (MC), somatosensory cortex (SC), retrosplenial cortex (RSC), association cortex (ASC), auditory cortex (AC), V1, and V2.

For urethane anaesthetized, acute experiments in Figure 4, mice were reared either ISO, ENR or normal (NOR) condition, in which three mice were reared with littermates in the standard cage after weaning at P19. All procedures involving animal care, surgery and sample preparation were approved by the Animal Experimental Committee of RIKEN Center for Brain Science, and performed in accordance with the guidelines of the Animal Experimental Committee of RIKEN Center for Brain Science.

### LFP recording

LFP recording was performed by fixing animals to a stereotaxic stage via a head frame. Silicon probes for chronic implanting (A1×16-5mm-50-177-HZ16_21mm with a TdT clip; NeuroNexus) were attached to a microdrive (dDrive-m; NeuroNexus). The microdrive was installed on the skull with dental cement to target the left hippocampus through a small craniotomy above CA1 (Bregma: mediolateral 1.8 mm, anteroposterior −1.8 mm). Electrode penetration was guided by monitoring the EEG waveforms and electrode depth. Mice were allowed to recover for at least one week before LFP recording. LFPs were recorded by an RZ2 multi-channel recording system at 24.4 kHz (Tucker-Davis Technologies, Alachua, FL, USA). For data analysis, LFPs were resampled at 20 kHz for SWR analysis and at 1250 Hz for theta analysis.

For acute experiments (Figure 4), a 16-channel linear silicon probe (inter-channel distance = 50 μm; Alx15-5 mim-50-177-A16; NeuroNexus, Ann Arbor, MI, USA) was used. Extracellular field potentials were recorded continuously with a sampling rate of 31 kHz (Digital Lynx; Neuralynx, Bozeman, MT, USA) (Shinohara et al., 2013; Tanaka et al., 2017). Body temperature was maintained at 37C throughout the surgery and recording sessions by a heat pad with rectal temperature feedback.

### Optical imaging

Mice were fixed to a stereotaxic stage by clamping the head frame and placed under a fluorescence stereomicroscope (MZ10F, Leica). The GFP3 filter set (excitation 470 ± 20 nm, emission 525 ± 25 nm, Leica) was used with an EL6000 light source (Leica). Images were acquired using ORCA-Flash 4.0 V3 CMOS cameras for chronic, and ORCA-Flash 4.0 V2 CMOS cameras for acute imaging, respectively. HC Image software (Hamamatsu Photonics) was used for image acquisition (25 Hz, 512 x 512 pixels, 16 bit) and shutter control. Electrophysiological recording was synchronized with imaging by feeding the 25 Hz image-acquisition TTL pulse.

### Simultaneous hippocampus and V1 EEG recording

We implanted electrode in the right hippocampus for SWR detection, an additional electrode in the left side of the visual cortex, and observed spike activities. V1 activity was filtered (250-750 Hz) and the resultant wave envelope was calculated and used for the detection of spiking activity with the following criteria: i) 10 SD larger than mean activities, and ii) the activity lasted longer than 1ms. Among the detected ripple activities, ripples with concurrent emergence of a sharp wave were confirmed visually and selected.

### Mouse state analysis

Mouse states were monitored by a digital camera during experimental sessions, and analyzed thereafter. Mouse states were classified as awake, drowsy, and sleeping. The awake state was defined as the presence of clear eye-opening and large pupils. Sometimes, the mice shook their bodies. In the sleep state, mice did not move and their eyes were firmly closed for more than 10s. States that were not clearly one of these two states were defined as the drowsy state and excluded from further analysis.

### Detection of brain states

EEG data analysis was carried out using MATLAB on computers running Linux or Windows.

#### SWR detection

Among simultaneously recorded LFPs from a silicon probe, the channel corresponding to the *stratum pyramidale* was identified by the presence of ripple oscillations. We found that electrode placement was stable as manifested by the channel location of ripples limited to within 100 μm (i.e., 2 channels of silicon probes) during chronic recordings over a week. For the analysis of SWR events, LFPs in the *stratum pyramidale* were first resampled to 20 kHz. Next, ripple events were detected automatically (29) with some improvements for noise reduction: abrupt changes in the high-frequency signal (80-250Hz; > 10 times SD) in multiple channels were removed as noise, and 96ms LPFs around noise were excluded from further analysis by zero-padding. The LFP was band-pass filtered for the ripple frequency band (125-250 Hz for chronic, and 80-250Hz for acute experiments) and the resultant signal was squared and then smoothed with a Hamming window of length 19.2 ms. In the first screening, ripples events were detected as the periods where the smoothed signal exceeded the mean value by 6.5 times the SD, with an inter-ripple interval of 100 ms. In the second screening, the local minima within ± 35 ms of each detected point (i.e., ripple trough) was assigned as the ripple timing and 200 ms ripple-filtered waveforms centered around the ripple timing were extracted for further analyses. After automatic detection, detected ripples were manually confirmed for the coincidence of sharp waves in the *stratum radiatum*.

#### Theta detection

The channel used for the detection of theta oscillations was located 200 μm below the *stratum pyramidale*. Theta oscillations were detected as described elsewhere (24, 29). We computed the EEG spectrogram in the CA1 *stratum radiatum* and identified periods that satisfied two criteria: (i) the ratio of the peak powers of the theta band (6.5–8.5 Hz) and the delta (2–3 Hz) band in each bin exceeded 0.6, and (ii) the period was at least 10s long. For acute experiment (Figure 4), we used the following criteria: (i) the ratio of the peak powers of the theta band (3.5–7.0 Hz) and the delta (2–3 Hz) band in each bin exceeded 0.6; (ii) the period was at least 15s long.

For theta phase analysis (Figure 5), we assigned the phase of the theta oscillations by approximating to a sine wave by Hilbert transform (in Figure 5A, the green dotted line shows the sine wave, and red and blue triangles indicate the peak and trough, respectively). Thereafter, we detected image frames nearest the trough, ascending, peak and descending phases, respectively.

### Imaging data analysis

For all imaging data, ΔF/F was defined by taking an averaged baseline as F. The baseline was calculated by removing astrocytic Ca2+ surges, which are typically (i) a rise of 8 % from baseline with (ii) slow dynamics that are synchronized across wide cortical areas (Monai et al., 2016). After the detection of SWR and theta events, images nearest the event were detected, and images during the sleep and awake states were selected according to the state of interest (i.e., awake or sleep). The calculated images for each session of the experiment were registered to a ‘standard image’ of the right hemisphere of the cerebral cortex, which represents all the cortical images, which allowed us to compare images between the experimental sessions and between mice. After image registration, we reduced image resolution by 2 x 2 binning.

### Analysis of imaging data around SWR events

We extracted 11 sequential images around SWR events; the SWR images were the seventh image in these sequences. The image sequences were named in terms of ΔF/Fs; i.e., −240ms, −200ms,…, +160ms, where the numbers indicate the time before or after SWRs. For the statistical significance map (Figures. 2A, 3B, and 4), the average intensities for the pixels in the cortical regions were calculated, and a t-test (against 0) was performed for each pixel. Pixels with p < 0.05 were visualized by the average ΔF/F values. The intensity transition graphs (Figures. 2B, 3B) were calculated by dividing the sum of intensities within a cortical area at the time of interest. Each cortical area was defined as shown in Figure 1B. For the bimodal index (Bim index) in Figure 2C, first, the areas surrounded by zero and intensities during phase 1 (−160ms to −40ms), and between zero and intensities during phase 2 (−40ms to 120ms) were calculated. The Bim index was calculated by dividing the difference between the phase 2 area and the phase 1 area (i.e., phase2 area – phase1 area) by the maximum intensity during phase 1. For the activation probability map (Figure 5A), the probabilities of positive changes of ΔF/F were calculated for each pixel, averaged over ISO animals, and mapped. For the quartile map, we first summed the positive signals ΔF/F for each SWR event as they occurred successively in the total brain area. After such successive calcium activities were summed, we selected the top quartile of the signals, and plotted the top quartile of calcium activities.

**Figure 2.**
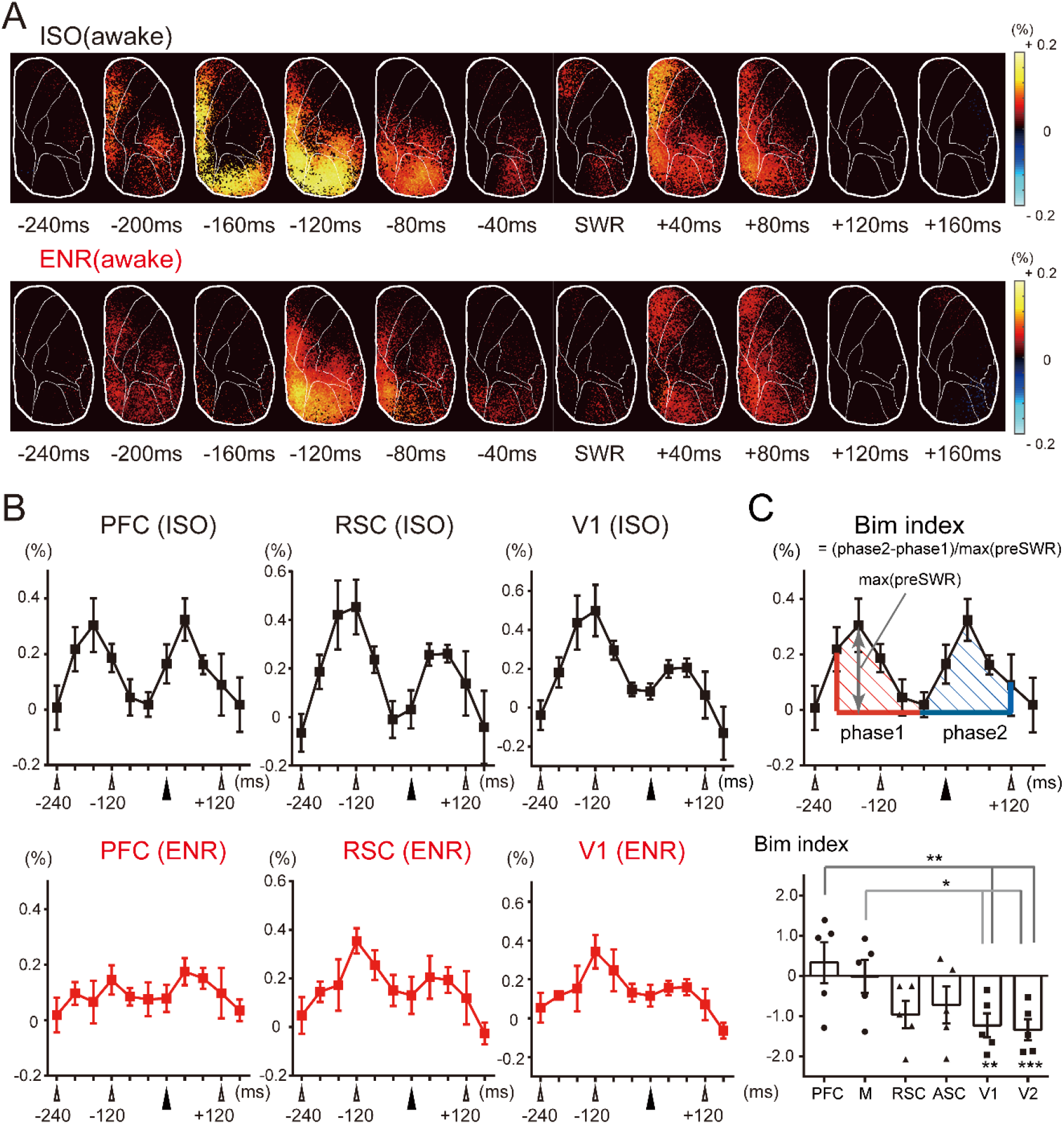
Cortical activity transition of awake mouse around SWR. (A) Upper row; Awake ISO mouse cortical activity transition from −240ms to +160ms around SWR (n =5) is shown as a percentage in color maps. In these maps, areas with a statistically significant increase or decrease from 0 (p < 0.05) are shown in the time lapse sequence. Lower row; Awake cortical activity transition maps around SWRs are shown for ENR mice (n =6). (B) Transitions of GCaMP ΔF/Fs signal from PFC, RSC, V1 are plotted for ISO mice (upper row) and ENR mice (lower row). The horizontal axis shows elapsed time (ms) with respect to SWR, and the vertical axis shows average ΔF/F values for specific cortical areas. Time of SWR is shown in the filled triangles. Error bars indicate SEM. (C) Upper column; Calculation of Bim index. The positive signal areas of Phase 1 (polygon made by −200ms --40ms lines; indicated by red diagonal lines) and Phase 2 (polygon made by −40ms – +120ms; indicated by blue diagonal lines) were calculated, and their difference was divided by the Phase 1 peak (indicated by the bidirectional arrow). A positive value of the Bim index means that the area of Phase 2 is greater than the area of Phase 1. Lower column; The Bim index was calculated for 6 cortical areas: PFC, MC, RSC, ASC, V1, and V2 in ISO mice. The statistical significance of differences was calculated between PFC and MC vs. V1 and V2, and between 0 versus V1 and V2. *p < 0.05; **p < 0.01; ***p < 0.001. Error bars indicate SEM.

**Figure 3.**
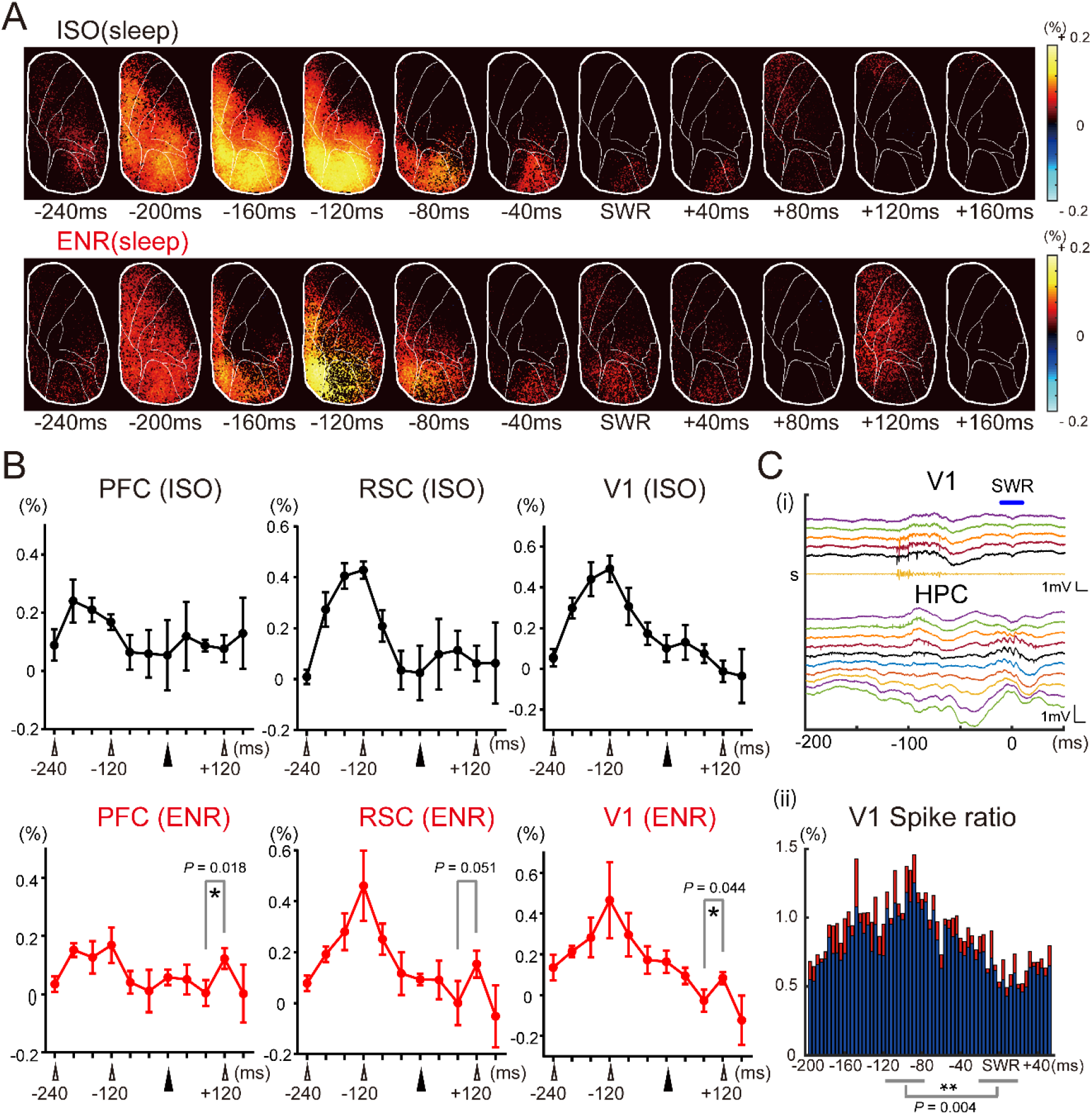
Cortical activity transition of sleep mouse around SWR. (A) Upper row; The cortical activity transition in natural sleep ISO mouse from −240ms to +160ms around SWR (n =5) is shown by percentages in color maps. In the maps, areas with a statistically significant increase or decrease from 0 (p < 0.05) are indicated in the time lapse sequence. Lower row; Sleep cortical activity transition maps observed in ENR mice (n =6). (B) Transitions of GCaMP ΔF/Fs obtained from PFC, RSC, and V1 are plotted for ISO (upper row) and ENR mice (lower row). The horizontal axis shows elapsed time (ms) with respect to SWR, and the vertical axis shows average ΔF/F values for specific cortical areas. Time of SWR is shown in filled triangles. In ENR mice, values at +80ms and +120ms were compared, and there were significant differences in PFC (p= 0.018; paired t-test) and V1 (p = 0.044). * p < 0.05. Error bars indicate SEM. (C) Simultaneous recording from V1 and hippocampus. (i) EEG traces from deep layer of V1, and hippocampus (HPC) were shown in upper and lower traces, respectively. The lowest trace in V1 (labeled as s) indicates V1 spike activities calculated from the 3rd lowest EEG trace in V1. The upper blue line indicates SWR. (ii) V1 spike rates from −200ms to +50ms to hippocampal SWR were calculated from 3 ISO mice. The spike rates between −200ms to +50ms were binned into 4ms time windows. Thereafter, the rates were normalized to sum value is 1, averaged over the animals, and indicated in the bar graphs. The average spike rates and ± SEM. of the spike rates are shown in blue and red bars, respectively. Statistically significant difference of spike rates was found between −120 to −80ms vs. −20 to 20ms (p = 0.004). ** p < 0.01.

**Figure 4.**
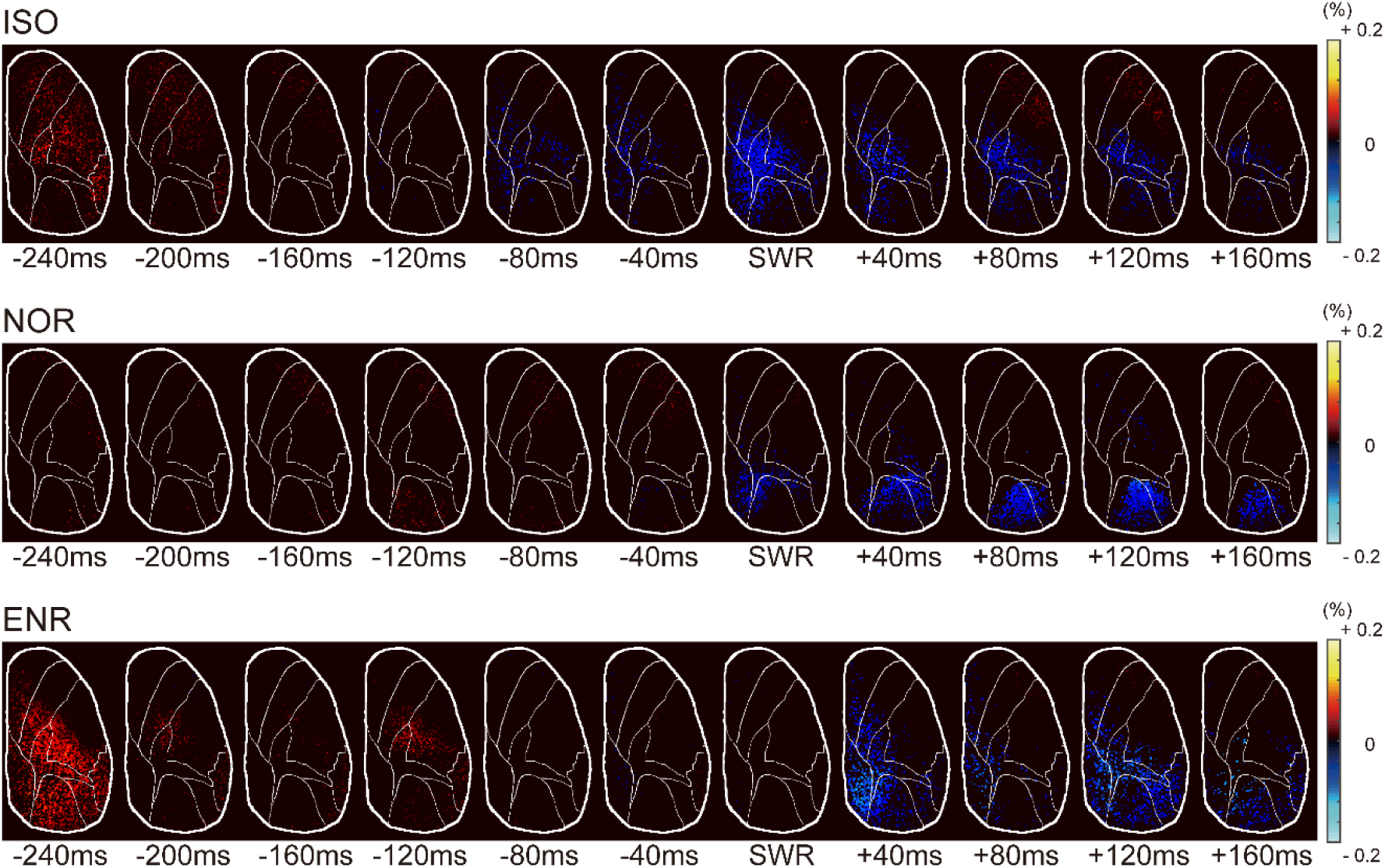
Cortical activity transition of urethane anesthetized mouse around SWR. Upper row; The cortical activity transition in urethane anesthetized ISO mouse from −240ms to +160ms around SWR (n =3) is shown by percentages in color maps. In the maps, areas with a statistically significant increase or decrease from 0 (p < 0.05) are indicated in the time lapse sequence. Middle row; Awake cortical activity transition maps around SWRs are shown for normal rearing condition mice (NOR) (n = 4). Lower row; Awake cortical activity transition maps around SWRs are shown for ENR (n = 4).

## Results

### Area-dependent bimodal cortical activation was observed around SWR in both ISO and ENR mice imaged during the awake condition

We imaged macroscopic cortical calcium activities on the right hemisphere of G-CaMP7 transgenic mice (Monai et al., 2016) (G7NG817) while simultaneously performing LFP recording in the contralateral hippocampal CA1 with a silicon multi-site probe. Two distinct hippocampal LFP states, theta and LIA, were detected based on their spectral property (see Methods). Since spontaneous cortical and hippocampal LFP events predominantly occur in both hemispheres (Tanaka et al., 2017; Vanni et al., 2017), this setting enabled investigation of the temporal dynamics between the hippocampus and the wide extent of the cortex. After weaning, transgenic mice were reared in either ISO or ENR condition for 4 weeks before recording (Figure 1A and B). Since calcium signals from both astrocytes and neurons are imaged in this transgenic strain, we confined our analysis to periods that did not contain astrocytic calcium events, thereby focusing on neuronal activities. We analyzed sleep and awake states separately. We first analyzed cortical activation during the awake LIA state. Cortical areas were defined as Figure 1C (Vanni et al., 2017), and the recording time and SWR numbers are shown in Table 1-1.

**Table 1-1.** Data of mice and recording time used for this study.

| **ISO mice** | NM016 | NM018 | NM019 | NM023 | NM030 |  |
| --- | --- | --- | --- | --- | --- | --- |
| ISO awake LIA | 2837.64 | 1112.8 | 5143.72 | 1060.72 | 1760.52 |  |
| Numbers of SWR | 462 | 362 | 1471 | 679 | 675 |  |
| Mean SWR interval | 8.1137 | 5.6664 | 4.5527 | 5.2280 | 4.1054 |  |
| ISO awake Theta | 1057.56 | 555.36 | 641.76 | 3283.44 | 1072.92 |  |
| ISO sleep LIA | 3131.24 | 3084.36 | 4221.64 | 5395.76 | 2804.64 |  |
| Numbers of SWR | 505 | 1156 | 919 | 2274 | 785 |  |
| Mean SWR interval | 6.5006 | 3.4413 | 4.7348 | 2.9594 | 3.7529 |  |
| ISO sleep Theta | 138.56 | 441.48 | 348.44 | 845.64 | 241.76 |  |
| Total recording time | 16800 | 14400 | 31200 | 26400 | 16800 |  |
| **ENR mice** | NM017 | NM020 | NM024 | NM025 | NM028 | NM032 |
| ENR awake LIA | 2616.4 | 6635.92 | 8765.84 | 11300.68 | 4345.36 | 1365.04 |
| Numbers of SWR | 523 | 901 | 913 | 3264 | 1183 | 335 |
| Mean SWR interval | 7.3819 | 7.7232 | 10.309 | 3.7362 | 4.1487 | 8.5062 |
| ENR awake Theta | 1011.64 | 1074.44 | 2207.76 | 966.04 | 962.2 | 2050.88 |
| ENR sleep LIA | 7065.4 | 6360.6 | 1535 | 6980.68 | 989.8 | 3739.68 |
| Numbers of SWR | 1916 | 879 | 252 | 2079 | 250 | 777 |
| Mean SWR interval | 4.8941 | 7.8449 | 6.3109 | 3.5089 | 3.4554 | 5.6408 |
| ENR sleep Theta | 1383.96 | 694.92 | 83.2 | 818.6 | 88.96 | 201.88 |
| Total recording time | 14400 | 28800 | 19200 | 60000 | 24000 | 38400 |

Five ISO and 6 ENR mice were used for the experiment. Total recording time, time in LIA state, and time in theta state used for analyses are presented in seconds. For LIA state, number of SWRs and SWR interval are also presented. Mean SWR interval is calculated by averaging SWR intervals after removal of long intervals (i.e., interval > 60s).

In ISO mice, cortical activation is observed in the medial part of the cortex including the PFC and RSC 200ms prior to SWR (abbreviated as −200ms). These increased activities diminished by −80ms in the PFC and −40ms in the RSC (Figure 2A; Table 2-1). Remarkably, a second activation peak was observed from 0 to +80ms in the PFC and from +40 to +80ms in the RSC. The activation in the posterior part of the cortex (V1 and V2) started at −160ms, and persisted until +80ms. In summary, the PFC exhibits a bimodal activation pattern that has peaks before and after SWR while the activities of V1 and V2 were observed relatively continuously around SWRs, with a mild sag at 0ms.

**Table 2-1.** Transition of cortical activity during awake condition.

|  | -240ms | -200ms | -160ms | -120ms | -80ms | -40ms | 0ms | +40ms | +80ms | +120ms | +160ms |
| --- | --- | --- | --- | --- | --- | --- | --- | --- | --- | --- | --- |
| V1 | - | + | ++++ | ++++ | ++ | - | - | + | ++ | - | - |
| V2 | + | +++ | +++ | ++++ | +++ | ++ | + | + | - | - | - |
| RSC | - | ++ | ++++ | ++++ | ++ | - | - | - | + | - | - |
| ASC | - | + | +++ | +++ | ++ | - | - | + | + | + | - |
| PFC | - | ++ | ++ | + | - | - | - | + | - | - | + |
| MC | - | + | ++ | + | - | - | + | ++ | + | - | - |
| SC | - | + | + | + | - | - | - | + | - | - | - |
| AC | - | + | ++ | ++ | + | - | - | + | - | - | - |

ENR awake
|  | -240ms | -200ms | -160ms | -120ms | -80ms | -40ms | 0ms | +40ms | +80ms | +120ms | +160ms |
| --- | --- | --- | --- | --- | --- | --- | --- | --- | --- | --- | --- |
| V1 | - | + | + | +++ | ++ | + | + | + | + | - | - |
| V2 | - | + | + | ++ | ++ | + | - | + | + | - | - |
| RSC | - | + | + | +++ | ++ | + | + | ++ | + | + | - |
| ASC | - | + | - | ++ | + | + | - | + | + | - | - |
| PFC | - | - | - | + | - | - | - | + | + | - | - |
| MC | - | - | - | + | - | - | - | + | + | - | - |
| SC | - | - | - | + | - | - | - | - | - | - | - |
| AC | - | - | - | + | + | - | - | - | - | - | - |

Transition of cortical activity shown in Figure 2B (awake states) is indicated in 5 grades by
symbols and colors: – (blank), + (green), ++ (yellow), +++ (orange), and ++++ (red), which
correspond to less than 0.1%, 0.1-0.2%, 0.2-0.3%,0.3-0.4%, and more than 0.4%,
respectively.

The time courses of activities from selected cortical areas are shown in Figure 2B. SWR-associated PFC activity exhibited a clear bimodal pattern, with first and second peaks of 0.304% at −160ms and 0.324% at +40ms, respectively. The RSC shows post-SWR milder activation (0.261% at +80ms) compared to pre-SWR (0.453% at −120ms). The activation curve of V1 shows a similar temporal change as RSC. Notably, the pre-SWR peak is higher than the post-SWR peak (0.500% at −120ms vs. 0.200% at +80ms). To evaluate the asymmetric nature of such bimodal activity between the pre-SWR and post-SWR periods, we introduced a “bimodal index” (Bim index), which indicates the magnitude of post-SWR activity relative to pre-SWR activity (Figure 2C; see Methods). The Bim index was calculated for representative brain areas in ISO mice: the value was 0.331 in the PFC and −0.0151 in the MC. The Bim index was −0.959 for the RSC, −0.721 for the ASC, −1.230 for V1 and −1.339 for V2. Statistically significant differences in the Bim index were seen for PFC vs. V1 (p = 0.00297) and MC vs. V1 (p = 0.0446). Moreover, the values for V1 and V2 were significantly below 0 (p = 0.00793 for V1 and p = 0.000980 for V2. On the other hand, the p value was 0.0756 for the RSC and 0.303 for the ASC.

In ENR mice, cortical activation also started at −200ms in the posterior part of the FC and RSC. Except for −160ms at which the cortical activities became less clear, the transition of the activated areas resembles to those observed in ISO condition, in lesser magnitudes of the activation intensities (Figure 2A, B, and Table 2-1). Though the Pre-SWR peak tended to be smaller than in ISO mice (ISO vs ENR Phase1 area (see Figure 2C): PFC, 0.653 vs 0.380, p = 0.055; RSC, 1.196 vs 0.930, p = 0.298; V1, 1.372 vs 0.867, p = 0.231), the PFC, RSC, and V1 all show bimodal activities similar to those in ISO (pre-& post-SWR peak intensities: RSC, 0.354% & 0.205%; PFC, 0.145% & 0.175%; V1, 0.343% & 0.159%). The magnitude of post-SWR peak relative to the first peak remained similar to that of ISO. Notably, V1 exhibited a more apparent pre-SWR single peak and a mild post-SWR peak. In all three cortical areas described above, the peak of the pre-SWR activation was observed at −120ms.

### Cortical activation was observed in both ISO and ENR mice during sleep, but predominantly in pre-SWR periods

Next, we asked if SWR-associated cortical activity during natural sleep resembles that of the awake state. As in Figure 2, we start with the analysis in ISO mice (Figure 3A). The pre-SWR activity peaked at −200ms in widespread middle and posterior cortical areas including the RSC, ASC and V1, V2, AC, and posterior PFC. Antero-lateral cortical areas did not have a prominent pre-SWR peak. The activation lasted until −120ms, and vision-associated areas such as RSC, V1, V2 and PFC exhibited markedly increased activities. At −80ms, PFC activation was diminished, and at −40ms only V1 and V2 remained active. After SWR, only the PFC exhibited weak activation from +80ms to +120ms.

In ENR mice, cortical activities started at −200ms, and the pattern resembled the ISO activity map. After −160ms, the activation areas were smaller than those of ISO and confined to the medial part of the cortex. From −80ms to SWR, the activated areas were larger than those of ISO, but the map shows more sparse patterns. Interestingly, at +120ms, we found a synchronized activation of sparse but pan-cortical areas, which was not observed in ISO (see below for quantification).

Figure 3B shows peri-SWR activities in the PFC, RSC, and V1 for ISO and ENR mice. Peri-SWR activity is characterized by a peak preceding SWR by 120-160ms. In ISO mice, the Bim index is −0.957 for the PFC, −1.94 for the RSC, and −2.06 for V1. In ENR mice, the respective values are −1.27, −1.97, and −1.78, respectively. As in the awake state, pre-SWR signal intensities of ISO mice tended to be higher although were greater than those in ENR, but this difference was not statistically significant (Phase 1 area; ISO vs ENR; PFC, 0.591 vs 0.417; p = 0.240 by t-test; RSC, 1.196 vs 1.148; p = 0.881; V1, 1.477 vs 1.239 p = 0.641). Moreover, ENR mice showed a brief post-SWR increase of activity between +80 and +120ms (+80-+120ms: PFC, 0.0385, 0.122, p = 0.0184; RSC, 0.0092, 1.53, p =0.0510; V1, −0.275, 0.844, p = 0.0439, paired t-test; Table 3-1.)

**Table 3-1.** Transition of cortical activity during sleep condition.

|  | -240ms | -200ms | -160ms | -120ms | -80ms | -40ms | 0ms | +40ms | +80ms | +120ms | +160ms |
| --- | --- | --- | --- | --- | --- | --- | --- | --- | --- | --- | --- |
| V1 | - | ++ | ++++ | ++++ | +++ | + | + | + | - | - | - |
| V2 | + | +++ | +++ | ++++ | +++ | ++ | + | + | - | - | - |
| RSC | - | ++ | ++++ | ++++ | ++ | - | - | - | + | - | - |
| ASC | - | + | +++ | +++ | ++ | - | - | + | + | + | - |
| PFC | - | + | + | + | - | - | - | - | - | + | - |
| MC | - | + | + | + | - | - | - | + | - | - | + |
| SC | - | + | + | + | - | - | - | - | - | - | - |
| AC | - | + | + | ++ | + | - | - | - | - | - | - |

ENR sleep
|  | -240ms | -200ms | -160ms | -120ms | -80ms | -40ms | 0ms | +40ms | +80ms | +120ms | +160ms |
| --- | --- | --- | --- | --- | --- | --- | --- | --- | --- | --- | --- |
| V1 | + | ++ | ++ | ++++ | ++ | + | + | - | - | - | - |
| V2 | + | ++ | ++ | +++ | ++ | + | + | - | - | - | - |
| RSC | + | + | ++ | ++++ | ++ | + | - | - | - | + | - |
| ASC | - | + | - | ++ | + | + | - | + | + | - | - |
| PFC | - | + | + | + | - | - | - | - | - | + | - |
| MC | - | + | - | + | - | - | - | - | - | + | - |
| SC | - | + | - | + | - | - | - | - | - | + | - |
| AC | - | + | + | ++ | + | - | - | - | - | - | - |

Transition of cortical activity shown in Figure 3B (sleep states) is indicated in 5 grades by
symbols and colors: – (blank), + (green), ++ (yellow), +++ (orange), and ++++ (red), which
correspond to less than 0.1%, 0.1-0.2%, 0.2-0.3%,0.3-0.4%, and more than 0.4%,
respectively.

We hypothesized that the bimodal peri-SWR activity pattern is caused by an increased SWR occurrence that will have multiple SWRs within the analyzed time period. However, the intervals between individual SWR events after removing long intervals (> 60s) tended to be longer in awake conditions (ISO awake vs. sleep; 5533.2ms vs 4277.8ms; p = 0.0656, ENR awake vs. sleep; 6967.5ms vs 5275.8ms; p = 0.0546; paired t-test; Table 1-1). Inter-SWR intervals did not differ significantly for SWR intervals shorter than 3000ms, either (Inter-SWR interval: ISO awake vs sleep, 707.1ms vs 691.1ms, p =0.284; ENR awake vs. sleep 732.2ms vs 764.7ms, p = 0.149). Therefore, SWR interval is not likely the cause of the differential post-SWR cortical activity pattern in ENR. Of note, we also analyzed urethane anesthetized mice (Karimi Abadchi et al., 2020) and found that results were distinct to natural sleeping mice regardless of rearing conditions (ISO, normal caged mice (NOR), and ENR; Figure 4). The results suggest that the distinct activity patterns between awake and sleep state are likely to be caused by natural awake/sleep cycles.

Our findings in the sleep condition can be summarized as follows. 1) Cortical activation precedes SWR period, especially in vision-associated areas. 2) Though cortical activation is mostly unimodal in ISO mice, post-SWR cortical activation of occurs at +120ms in ENR mice. 3) Peri-SWR activation in ISO mice is confined to the frontal and motor areas, whereas it is more widespread in ENR mice.

As peri-SWR cortical activities show a pronounced pre-SWR activation, we further sought to verify this by unit recordings in V1 during sleep (Figure 3C). We observed SWR-associated unit activities in the visual cortex during sleep (n = 3 ISO mice: 168, 199, and 338 SWRs). The SWR-triggered averaging of V1 multi-unit activity shows a broad elevation of activity with a peak between −120ms and −80ms. For instance, normalized V1 spike activity from −120ms to −80ms is significantly higher than that from −20ms to +20ms (10.9% vs 5.8%, multiunit activity normalized to the −200 to +50ms period, p = 0.00441; t-test). These data are consistent with the calcium imaging data.

### Greater calcium elevations tend to be observed during post-SWR periods, and both the animal state and rearing conditions modulate the occurrence of peri-SWR cortical activation timing

Given the awake/sleep state-dependent bimodal and unimodal peri-SWR cortical activation, we sought to investigate how the animal’s state brings about differential activation. Since ISO mice clearly exhibit this differential activation pattern, we first focused on ISO animals. We asked if the probability of cortical activation (i.e., emergence of positive values of ΔF/F) that accompanies SWRs is similar among individual SWRs or varies across each event. We calculated the probability of calcium elevations regardless of the magnitude, and plotted the activation probability maps for awake and sleep periods (Figure 5A). During pre-SWR periods, pixels with > 50% activation probability were found in the posterior 2/3 of the cortex in both the awake and sleep states throughout the −200ms to −80ms period. In the central part of the cortex, the probability of pre-SWR activity is higher in the awake state. By contrast, post-SWR activity becomes higher in peripheral cortical areas in awake states (0 ms to +80 ms). The result indicates that the probabilities of the calcium elevation during post-SWR periods are higher in awake animals.

**Figure 5.**
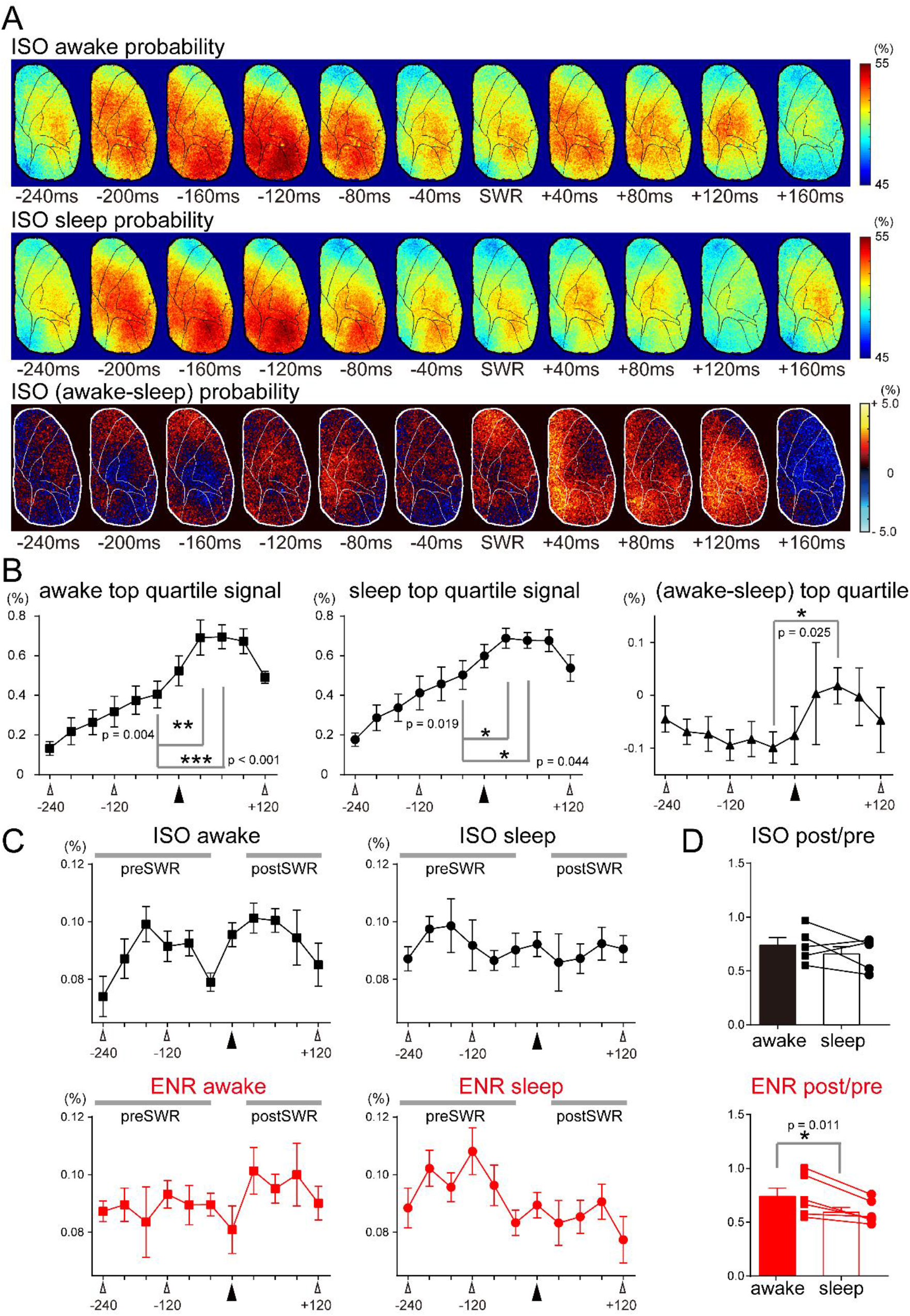
Distinct cortical activities between awake and sleep state around SWR. (A) Probability maps of cortical activation in ISO mice associated with individual SWRs, averaged across 5 ISO mice. Upper row: Probability maps of cortical activation in the awake condition. Middle row: Probability maps of cortical activation in the sleep condition Lower row: Probability maps of the difference in cortical activation between the awake and sleep conditions. (B) Top quartile of positive cortical signals of ISO mice are plotted for awake (left panel) and sleep (middle panel) states, and their difference (right panel). Time of SWR is shown in filled triangles. * p < 0.05; ** p < 0.01; *** p < 0.001. Error bars indicate SEM. (C) Distribution of the time of maximum positive signal intensities around SWR associated cortical activation. Time of SWR is shown in filled triangles. Upper panel: left, ISO awake; right, ISO sleep. Lower panel: left, ENR awake; right, ENR sleep. (D) Comparison of the distribution of maximum signals between pre-SWR and post-SWR periods. Post/pre-SWR ratios of maximum signals were calculated between ISO (upper) and ENR (lower panel). * p < 0.05. Error bars indicate SEM.

Next, we analyzed the magnitude of the calcium events. The cortical calcium signal transitions shown in Figures. 2 and 3 are the averages of total events which include both increases and decreases of ΔF/F (Figure 5-1). We hypothesized that individual cortical activations during the post-SWR period are larger in the awake state than in the sleep state. The top quartiles of the positive activity of ΔF/F were plotted for awake and sleep states (Figure 5B). We found that top quartile activity peaks predominantly in post-SWR periods in both states. The increase in the quartile signal from −40ms to +40ms is 70.1% (p = 0.00398; paired t-test) and that from −40ms to +80ms is 71.8% (p < 0.001) in the awake condition. These increases were 36.8% (p = 0.0193) and 34.5% (p = 0.0435), respectively, in the sleep condition. Thus, the increase of cortical activities around SWR are larger in the awake condition. Accordingly, the difference between the awake and sleep quartiles increases around SWR, and, although the difference between −40ms (−0.984%) and 40ms (0.328%) is not significant, that between −40ms and +80ms (0.184%) is statistically significant (p = 0.0254). In conclusion, diphasic and monophasic differences between awake and sleep conditions can be partially ascribed to the top quartile of positive cortical activities after SWR.

**Figure 5-1.**
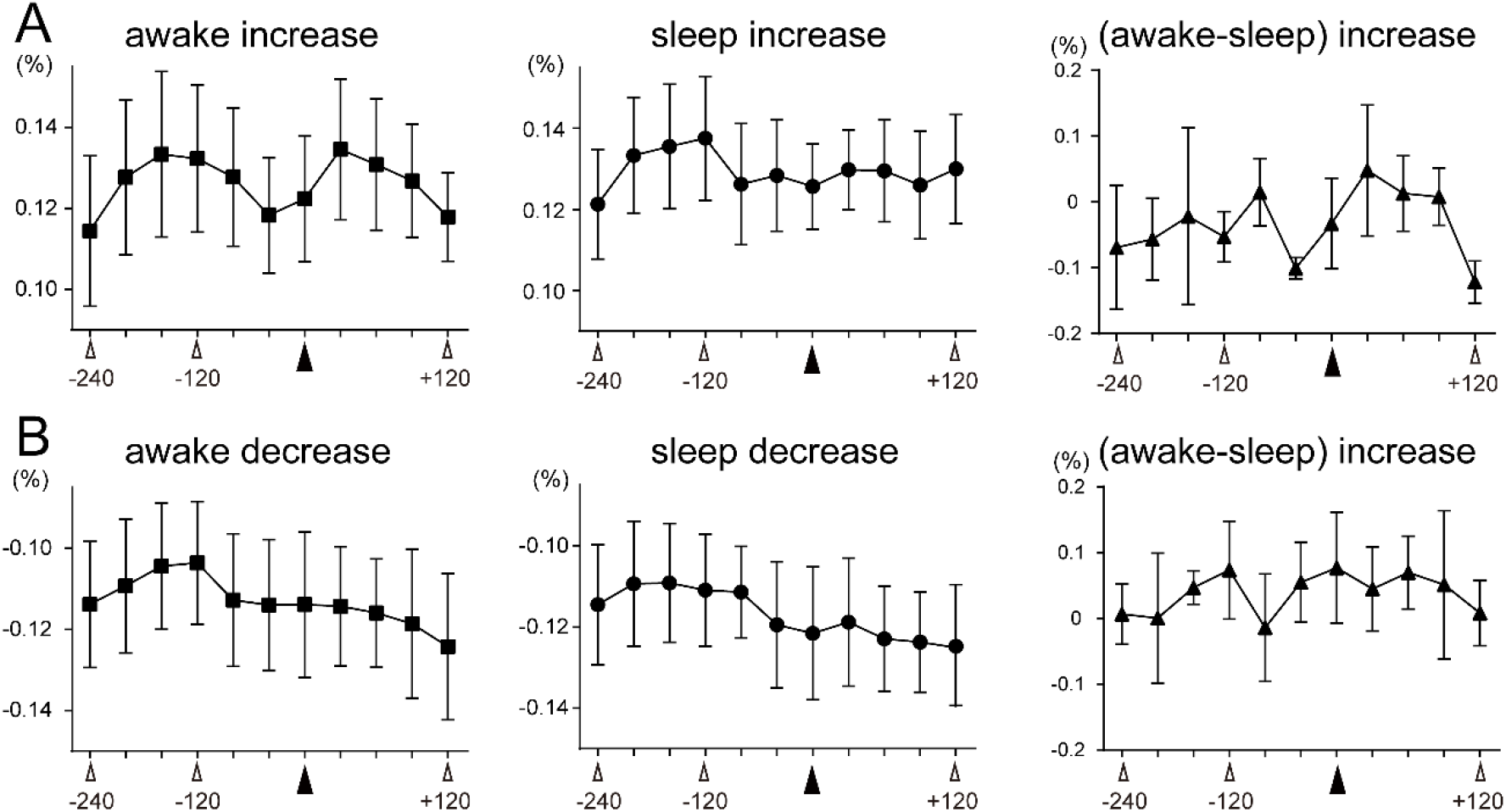
Increase and decrease of calcium activities around SWR obtained from the whole cortical areas of ISO mice. Increase (A) and decrease (B) of signal throughout the cortical areas calculated from ISO animals. Left, middle, and right panels show the time course of awake, sleep and the difference between awake and sleep signal intensities. Time of SWR is shown in filled triangles. Error bars indicate SEM.

The data in Figure 5-1 also indicates that the decreases of ΔF/F are considerably large (on average, 90.0% and 90.4% of increases in the awake and sleep conditions, respectively), which strongly implies that SWR-associated cortical activation is not a simple process that is accompanied by hippocampal SWRs. Next, we plotted the time at which each cortical SWR-triggered trace exhibited maximum intensity (Figure 5C). These times exhibited a dispersed distribution. Interestingly, ISO mice in the awake condition exhibited a bimodal distribution, while ISO mice in the sleep condition showed a monotonous distribution. In ENR mice, the distribution of maximum intensities in awake and sleep mice are mildly concentrated during the post-SWR and pre-SWR periods, respectively. The ratios of pre-vs. post-SWR (post-SWR/pre-SWR) were calculated and compared between the awake and sleep states; the values are 0.739 vs. 0.567 (p = 0.363; paired t-test) for ISO, and 0.740 vs. 0.591 (p =0.0110; paired t-test) for ENR (Figure 5D). Even though bimodal activation in ENR is less clear than that in ISO (Figure 2B), the results indicate a similar temporal bias of SWR-associated cortical activation to ISO condition.

### State-dependent cortical dynamics during the theta period are modulated by environmental stimuli

So far, we have investigated the spatio-temporal coordination of cortical activity relative to hippocampal SWRs and described the state-dependent activity patterns. Next, we analyzed cortical calcium dynamics during theta oscillations. First, we compared cortical calcium signal intensities between SWR and theta periods. Theta periods were detected from an EEG trace in the *stratum radiatum* of CA1 (e.g., Figure 6A), and cortical fluorescent images during the periods were compared with cortical activities close to SWRs. In the awake condition, calcium activity during theta periods was stronger in the medial half of the cortex in ISO mice, but a difference was not detected in ENR animals (Figure 6B). During the sleep state, no areas exhibited a significant difference between theta and SWR in ISO and ENR mice.

**Figure 6.**
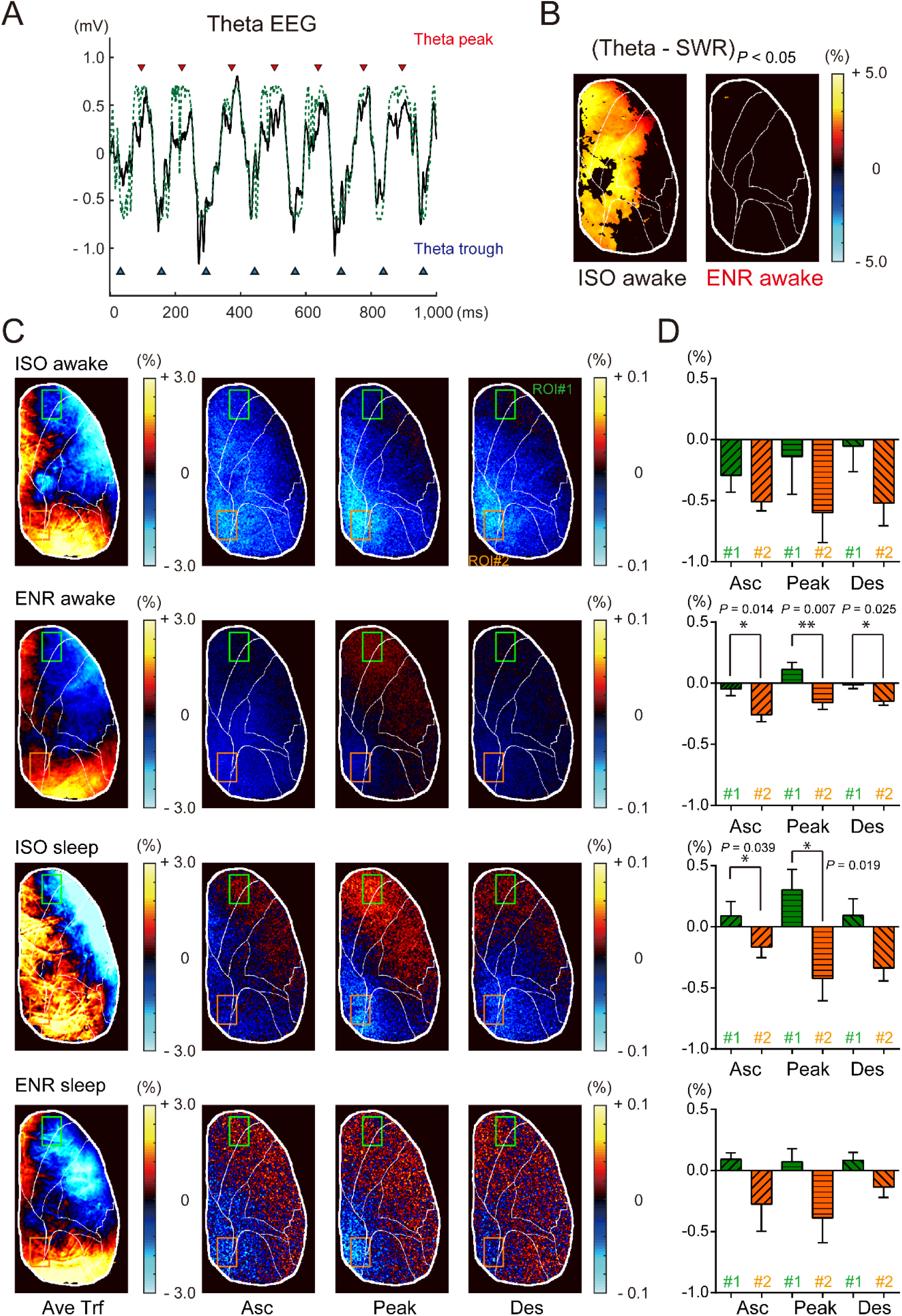
Cortical activities during hippocampal theta state. (A) An example of theta oscillations (~ 8Hz) recorded from the CA1 *stratum radiatum* (200 μm below the *stratum pyramidale)* of an ISO sleep mouse (black line). Dotted green line indicates EEG trace approximated to a sine waveform by Hilbert transformation. Red and blue triangles above and below the EEG trace indicate theta peaks and troughs, respectively. (B) Maps of statistically significant areas (p < 0.05) of (Theta – SWR) intensities in ISO (left panel) and ENR (right panel) mice. In ISO mice, a statistically significant cortical area spread from the midline to nearly half of the medial side of the cortex. In ENR mice, the differences in intensities are mostly diminished. (C) Left panel: Maps of calcium activities during the theta trough phase averaged across all of the cortical areas (Ave Trf). Maps of ISO awake, ENR awake, ISO sleep, and ENR sleep mice are shown in this order from top to bottom. In all of the experimental conditions and states, the antero-medial and posterior parts of the cortex show higher activities than the antero-lateral part of the cortex. Rectangles surrounded by green and orange lines are ROI#1 and ROI #2, respectively. Right panels: Differential activity map of the ascending (Asc; left map), peak (Peak, middle map) and descending (Des; right map) phases, which correspond to the left panel. (D) Bar graphs of differential intensity changes in ROI#1 and ROI#2 intensities from the trough phase. *p < 0.05; **p<0.01. Error bars indicate SEM.

Next, we asked if the hippocampal theta rhythms are correlated with cortical activity. For each frame of cortical calcium imaging, we computed the corresponding phase of theta oscillations (Figure 6A, see Methods). Thereafter, we computed the averaged image from images captured at the trough phase (Figure 6C). Interestingly, there was a spatial gradient in signal intensity, which varied by both animal state and rearing condition. In all conditions, the posterior cortex and anterior half of the cortex in the midline manifested higher activities, whereas the antero-lateral part of the cortex exhibited lower activities. Except for ISO mice in the sleep state, black areas, which indicate cortical areas with average activity, are L-shaped (Figure 6C), the frontal, RS and visual cortex have higher activities, and the motor and sensory cortex have lower activities. Surprisingly, the cortical activity maps of ISO awake and ENR sleep appeared similar. Further, we examined whether cortical activities show oscillatory activities with hippocampal theta rhythm. The relative change to the trough phase is computed for ascending, peak, and descending phases (Figure 6C). Except for anterolateral part of the cortex in ISO awake condition, the change in intensity in the peak phase is in the direction opposite the averaged activities (i.e., increase in the anterolateral part and decrease in the postero-medial part of the cortex). The result indicates that, differences in calcium activities across cortical areas are highest in the trough phase and reduced in the peak phase.

Since the extreme anterior and posterior parts of the cortex exhibited marked differences between the trough and peak phases, we set ROI#1 and ROI#2 in the extreme frontal and postero-medial parts of the cortex, respectively, and plotted the changes in intensity. In awake mice, subtraction images were negative throughout almost the entire cortical area expect for the anterior half in the peak phase of ENR mice. Thus, cortical activities are highest in the trough phase. During the peak phase of awake ENR mice, ROI#1 exhibited a positive direction of change. During the ascending, peak and descending phases, ROI#1 and ROI#2 exhibited significant differences in changes in activity (p = 0.0137, 0.00671, and 0.0249, respectively). In sharp contrast to the awake condition, ROI#1 and ROI#2 exhibited opposing changes in sleep mice (Figure 6D). The trend is apparent in ISO mice, and the difference from the trough phase was significant for the ascending and peak phases (p = 0.0389 and p = 0.0190). ENR mice exhibited similar tendencies, but the differences were not significant. Although the average intensity maps of theta were similar between ISO awake and ENR sleep mice, their oscillatory behaviors are distinct, and their oscillatory dynamics basically depend on the brain state.

## Discussion

In this study, we combined transcranial cortical calcium imaging with hippocampal LFP recording in unanesthetized mice and addressed the spatial extent of cortical dynamics with respect to hippocampal activity. High-speed calcium imaging leads to the identification of cortical areas that are associated with hippocampal SWRs. Vision-related areas such as the visual and RS cortex are highly associated with hippocampal SWRs. Furthermore, cortical activity relative to hippocampal SWR depended on brain awake/sleep state. We also unveiled the phase-locked pan-cortical activity to hippocampal theta oscillations.

We found a widespread cortical activity occurs shortly before (−240ms to −80ms) hippocampal SWR. While such an activity pattern is pronounced in vision-related areas, visual inputs unlikely the primary drive since this pattern is present during sleep mice. The data suggest that cortical activities in various areas are involved in triggering hippocampal SWRs (Sirota et al., 2003; Rothschild et al., 2017). Surprisingly, ENR mice exhibited moderate elevation compared to ISO mice. The finding that ENR mice showed less consistent cortical activation than ISO mice might be because ENR mice have more variations between individual subjects (Korholz et al., 2018).

In sharp contrast to the pre-SWR period, we identified distinct post-SWR cortical activity patterns depending on sleep/awake state. Considering the widely acknowledged notion of hippocampal replay of recently acquired experience (Wierzynski et al., 2009; Carr et al., 2011; Silva et al., 2015), the relatively lower post-SWR cortical activity in the sleep state is unexpected. Though Bim index varies depending on the cortical area, our analysis indicates that SWR-associated cortico-hippocampal information flow is generally unidirectional during sleep period in that cortical activity precedes SWR. On the contrary, cortical areas associated with vision (V1, V2, RSC and PFC) exhibits post-SWR activity in the awake state hinting at a closed loop of cortex-hippocampus-cortex.

The entorhinal cortex and hippocampus are reciprocally connected and physiologically tightly linked (Witter, 1993; Chrobak et al., 2000; Moser et al., 2014). Our results indicate that widespread cortical regions might be involved in a bidirectional excitation loop on a longer time-scale. The proportion of bidirectional activities depends on the cortical area, and the frontal part of the cortex has a relatively higher tendency for activation in the post-SWR period compared to pre-SWR activation. Accordingly, the Bim index for awake conditions is arranged in a spatially organize manner (Figure 7). Specifically, cortical areas are grouped in the following order: i) PFC and MC, ii) RSC and ASC, and SC iii) V1 and V2 and AC. Interestingly, the three areas are arranged in the anterior-posterior axis of the cortex, and the intensities of cortical activation during pre-SWR tend to be in the opposite order. This finding is indicative of information transfer from the hippocampus to the cortex via SWR and integration to sensory-motor processing, in line with previous work that suggested a role of hippocampal SWR in immediate future spatial navigation (Jadhav et al., 2012; Pfeiffer and Foster, 2013).

**Figure 7.**
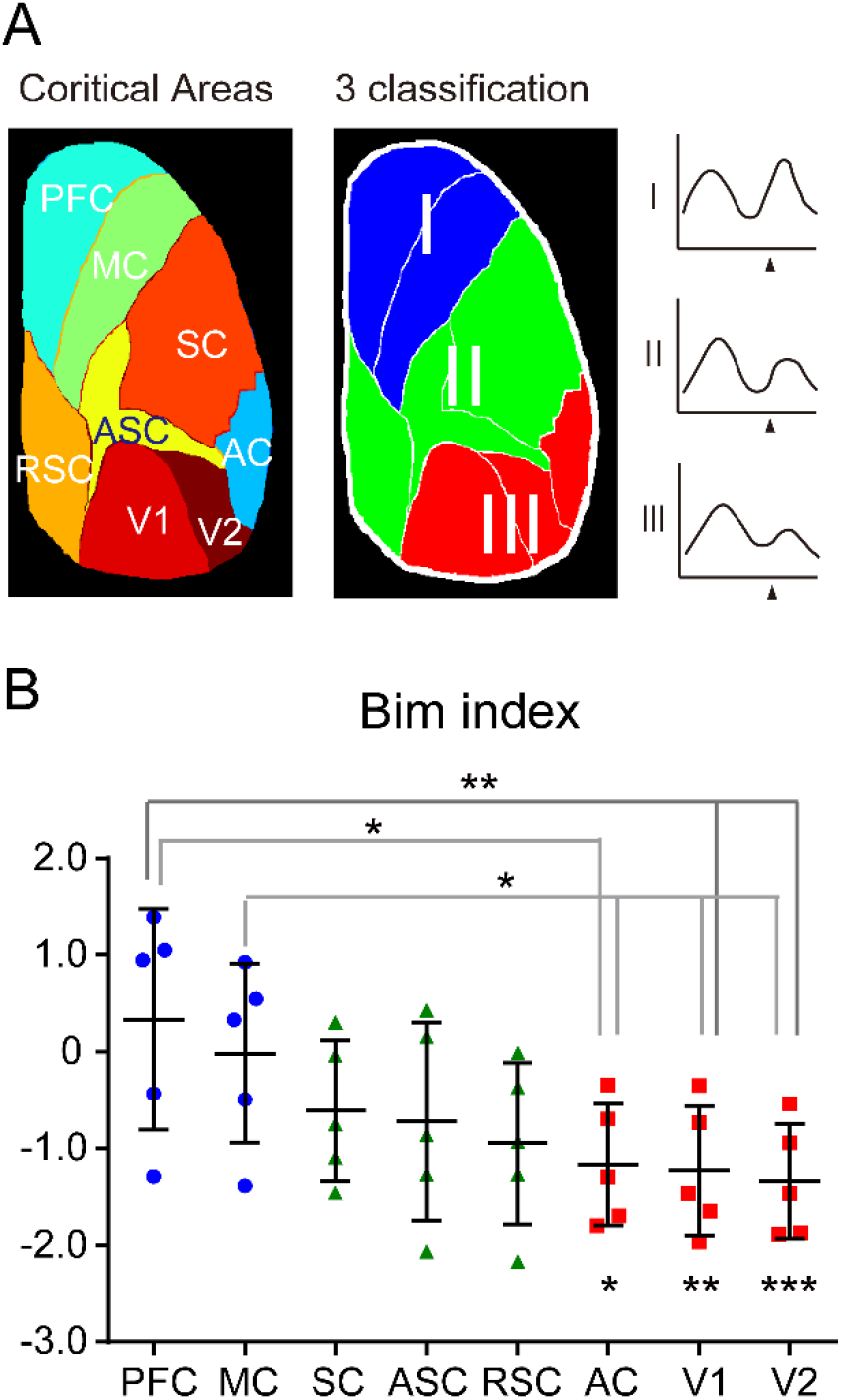
Cortical area classification by Bim index during awake period. (A) Cortical areas (left column), and 3 classification (Region I, II, III) of the areas (right column) defined by Bim index during awake period of ISO mice. Calcium elevation of Region I (PFC and MC) shows clear bimodal pattern, whereas that of Region III (V1 and V2 and AC) exhibits nearly monophasic dynamics during peri-SWR period. Region II (SC, ASC, and RSC) show medium pattern of elevation between Region I and Region III. (B) Bim index (Figure 2C) of all cortical areas. There are significant differences between Bim index of areas in Category I and Category III. Significant differences were also found between Bim index of areas included in Category III and 0. *p < 0.05; **p < 0.01; ***p < 0.001. Error bars indicate SEM.

During the post-SWR period, sleep ENR mice exhibited low cortical activities, similar to ISO mice, except at +120ms, whereby synchronized surge of pan-cortical activities emerged (Figure 3B). This observation strongly implies that environmental stimuli induce the integration of various modalities processed in the cortex by promoting coherent activities of cortico-cortical networks (Maviel et al., 2004). While the modulation of cortico-hippocampal dynamics after memory acquisition is outside the scope of this paper, examination of post-SWR co-activation of various cortical areas will be valuable (Battaglia et al., 2004; Wang and Ikemoto, 2016).

Investigation of individual SWRs revealed that SWR-associated cortical activity is broadly distributed and its spatial pattern varies at each SWR event. The bimodal distribution in the awake state is possibly due to multiple factors. Both probabilities and magnitudes of the cortical activity are larger in awake condition than sleep during post-SWR period in ISO mice (Figure 5A). Smaller calcium events frequently occur in the pre-SWR period, while greater events happen less frequently in the post-SWR period (Figure 5B). In addition, modulation of timing of maximum cortical activities around SWRs are modulated by animal status in both ISO and ENR mice (Figure 5C and D). Interestingly, the dynamics in ENR mice tended to show a skewed distribution, with clustering post- and pre-SWR, respectively, depending on the animal status. As ENR mice receive more sensory stimuli than ISO mice, this shift might be caused by brain state-dependent switching of cortico-hippocampal information flow. After SWRs, the cortex of ENR awake mice might be processing more abundant information that is associated with spatial information processed in the hippocampus.

Finally, theta-phase modulation of cortical activities was mapped for the first time at the cortex-wide level at submillimeter resolution. While theta phase-locked activity between the PFC and hippocampus has been well-established for spatial navigation (Siapas and Wilson, 1998; Jones and Wilson, 2005), our data indicate that theta phase-locked activity is spread across the entire cortex and advocate that phase-dependent information coding in humans (Kunz et al., 2019) is preserved in mice. Interestingly, although higher cortical activities in the posterior and medial part of the cortex than antero-lateral part were commonly observed in both awake and sleep conditions, (Figure 6C, trough maps), the dynamics with respect to hippocampal oscillatory activities are dependent on the brain state. Distinct phase-locked activities in memory encoding vs. retrieval (Hasselmo et al., 2002; Kragel et al., 2020) might be one of the reasons of differential state-dependent behaviors. In conclusion, in both the LIA and theta states, the brain state primarily regulates cortico-hippocampal dynamics, and the postnatal environment significantly impacts on their communication.

## Conflict of Interest

The authors declare no competing financial interests.

## Acknowledgments

This work was supported by KAKENHI 17H02221, 26282222, 16H06404, 16H01888. We are grateful for valuable discussions with Reiji Yamazaki and Nobuhiko Ohno.

## Author Contributions

YS and HH conceptualized the study. KS and YS performed the experiments. YS performed the data analysis. YS and HH wrote the manuscript. TU supervised the study. TU and YS acquired funding.

## Notes

### Competing Interest Statement

The authors have declared no competing interest.

